## Supplementary figures and images for "Zika virus infection drives epigenetic modulation of immunity by the histone acetyltransferase CBP of *Aedes aegypti*"

### Supplementary Figure 1

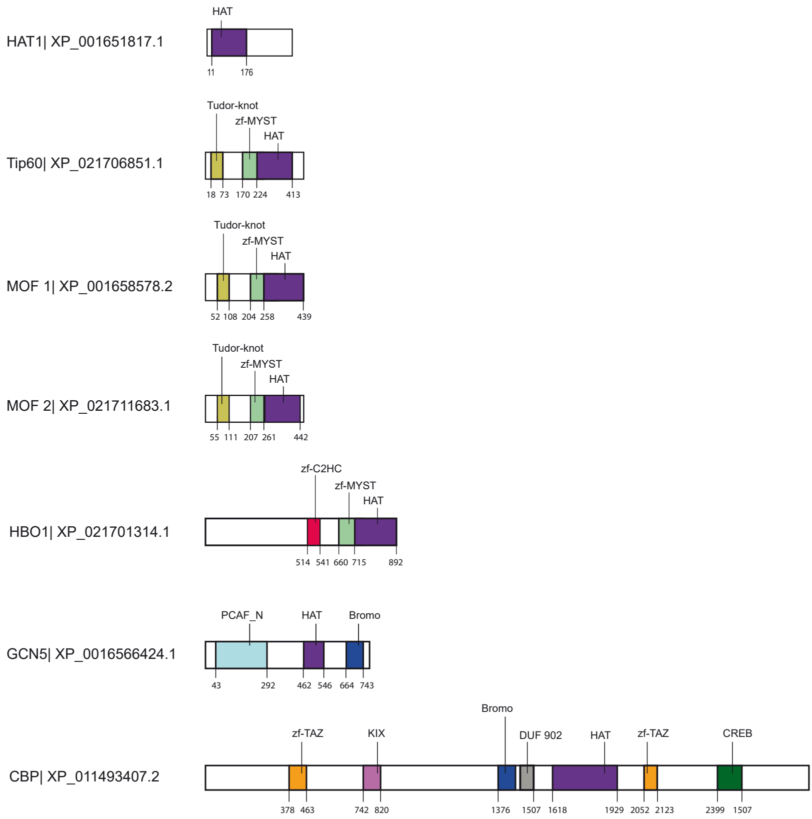

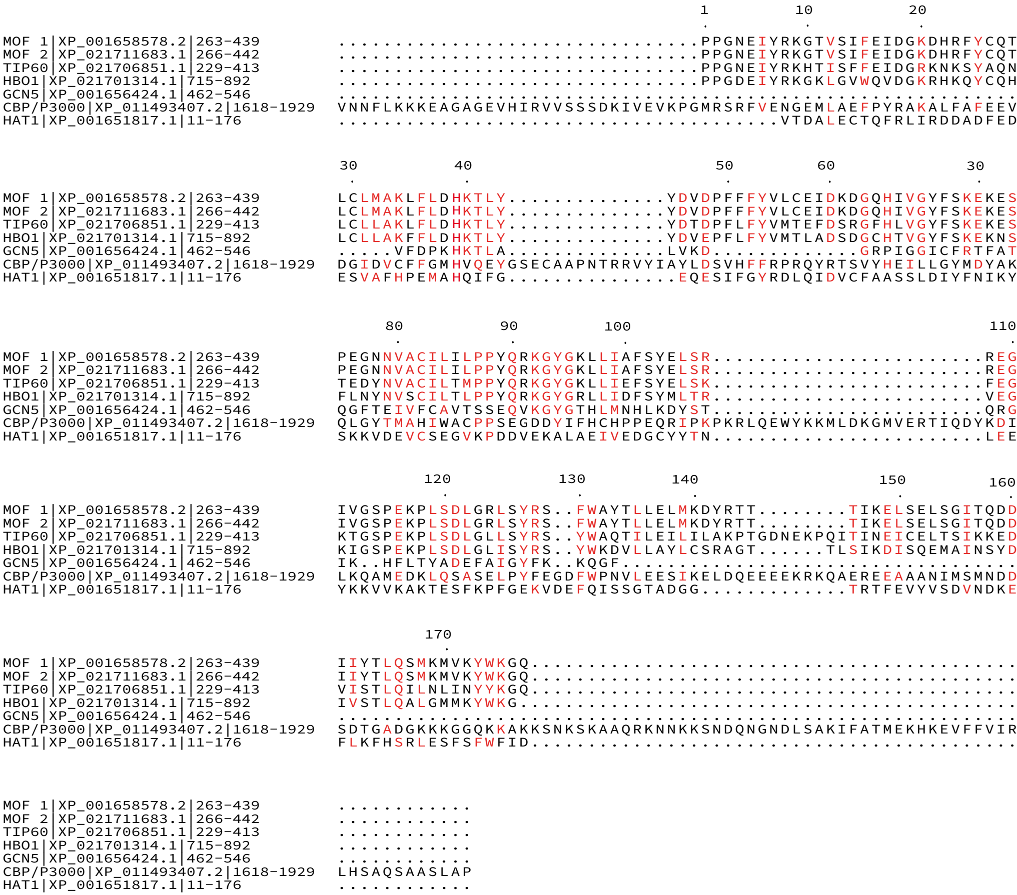

### Supplementary Figure 2

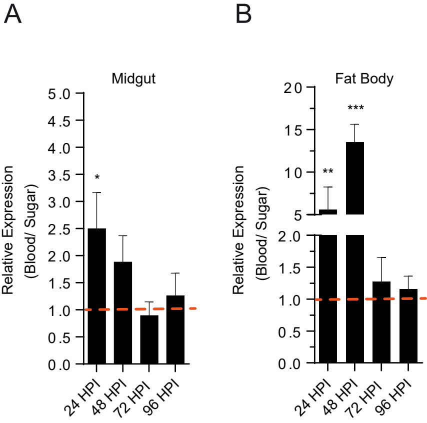

### Supplementary Figure 3

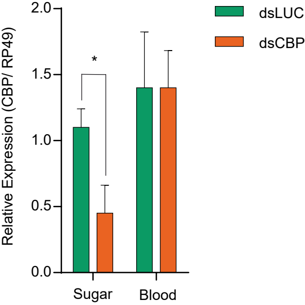

### Supplementary Figure 4

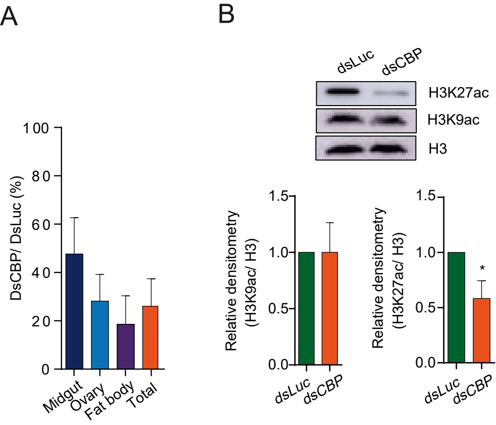

### Supplementary Figure 5

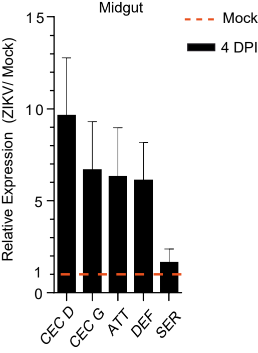

### Supplementary Figure 6

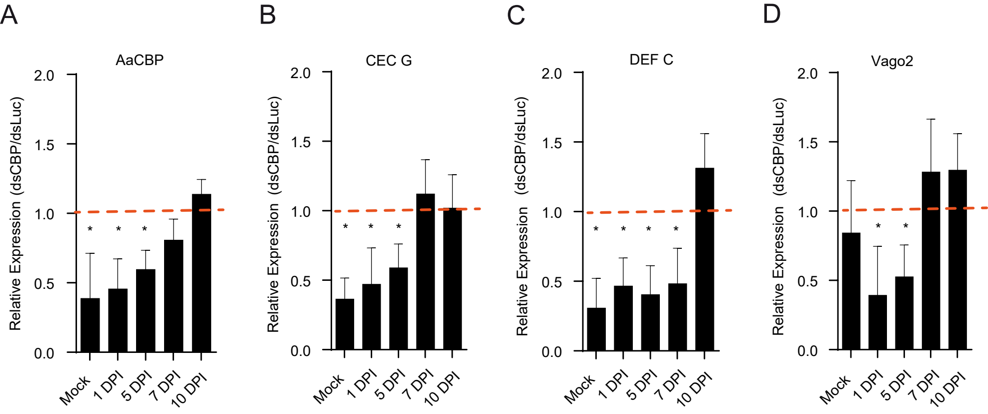

### Supplementary Figure 7

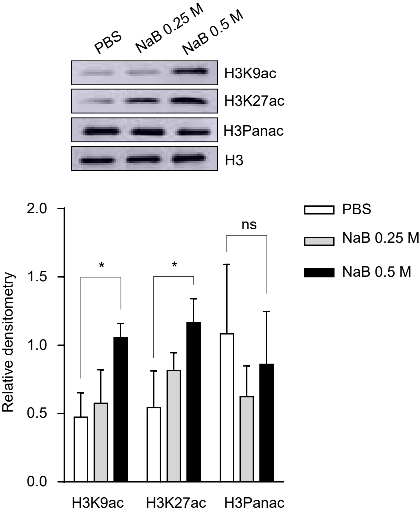
