## Supplementary Table 1 for "Zika virus infection drives epigenetic modulation of immunity by the histone acetyltransferase CBP of *Aedes aegypti*"

Supplementary Table 1. List of primers used in this study

| Primer use | ID VectorBase | Primer name | Sequence (5'-3') |
|---|---|---|---|
| qPCR | AAEL013816 | AaCBP Forward | CTCTCGATACTGAAAGCCAACC |
| | | AaCBP Reverse | TGCTGCATTACGTTGTGAGG |
| | AAEL003396 | RP49 Forward | GCTATGACAAGCTTGCCCCCA |
| | | RP49 Reverse | TCATCAGCACCTCCAGCTC |
| | AAEL029041 | Cecropin D Forward | GAAGAAGCTGGGAAAGAAATTG |
| | | Cecropin D Reverse | CCAATCGCTTTTATTCCTACAAC |
| | AAEL029038 | Cecropin G Forward | GTTATTTCTCCTGATCGCCG |
| | | Cecropin G Reverse | CTCGTTTTCCTGCACCTCCC |
| | AAEL003832 | Defensin C Forward | CTTTGTTTGATGAACTTCCGGAG |
| | | Defensin C Reverse | GAACCCACTCAGCAGATCGC |
| | AAEL003841 | Defensin A Forward | CTATCAGGCTGCCGTGGAG |
| | | Defensin A Reverse | CAATGAGCACAAGCACTATC |
| | AAEL003389 | Attacin Forward | TTGGCAGGCACGAATGTCTTG |
| | | Attacin Reverse | TGTTGTCGGGCACGGGAAGTG |
| | AAEL014078 | Serpin Forward | ACGTGATGGATTGGATGGAG |
| | | Serpin Reverse | GTGCCTGCACTTGTTTCTGA |
| | AAEL006794 | Dicer 2 Forward | CACCGACACAGTTAGCAAGT |
| | | Dicer 2 Reverse | CCATCTTCCGGTCTGATTCTTC |
| | AAEL017251 | Agonauta 2 Forward | TACCCGGCTCCAACCTATTA |
| | | Agonauta 2 Reverse | CTGCATTCTCTCGTACTCCTTG |
| | AAEL000165 | Vago 2 Forward | CCGGTGAATGCTACGATACTGA |
| | | Vago 2 Reverse | TGAGAAATCCGTTCCACATGAC |
| dsRNA | AAEL013816 | dsAaCBP1 Forward | TAATACGACTCACTACTATAGGGAGAGTTAG GCCTCAATCAGCTACTC |
| | | dsAaCBP1 Reverse | TAATACGACTCACTACTATAGGGAGAGAGGT AAACAAACCGGACAG |
| | | dsAaCBP2 Forward | TAATACGACTCACTACTATAGGGAGACGTTA ACGTTCTCTCCCAAC |
| | | dsAaCBP2 Reverse | TAATACGACTCACTACTATAGGGAGATTGGA TAGCTGATCCTGTGC |
|  | ----- | dsLuc Forward | TAATACGACTCACTACTATAGGGAGACTGGA GACATAGCTTACTG |
| | | dsLuc Reverse | TAATACGACTCACTACTATAGGGAGAGGATC TCTCTGATTTTTCTTGCG |
